# Presence of Atopy Increases the Risk of Asthma Relapse

**DOI:** 10.1101/217547

**Authors:** Laurel Teoh, Ian Mackay, Peter P Van Asperen, Jason Acworth, Mark Hurwitz, John W Upham, Weng Hou Siew, Claire YT Wang, Theo P Sloots, Teresa Neeman, Anne B Chang

**Author notes:** Address for correspondence: Dr Laurel Teoh, Department of Paediatrics and Child Health, Centenary Hospital for Women and Children, PO Box 11, Woden, ACT 2606, Australia. Deceased 19 November 2015 (Peter P Van Asperen).

## Abstract

**Objectives:** In children with hospitalised and non-hospitalised asthma exacerbations, to: (a) describe the point prevalence of respiratory viruses/atypical bacteria using polymerase chain reaction (PCR) and; (b) evaluate the impact of respiratory viruses/atypical bacteria and atopy on acute severity and clinical recovery.

**Design:** This was a prospective study performed during 2009-2011.

**Setting:** The study was performed in the Emergency Departments of 2 hospitals.

**Patients:** 244 children aged 2-16 years presenting with acute asthma to the Emergency Departments were recruited. A nasopharyngeal aspirate and allergen skin prick test were performed.

**Main outcome measures:** The outcomes were divided into (a) acute severity outcomes [Australian National Asthma Council assessment, hospitalisation, Functional Severity Scale, acute asthma score, asthma quality of life questionnaires for parents (PACQLQ) on presentation, asthma diary scores (ADS) on presentation and length of hospitalisation] and (b) recovery outcomes (PACQLQ for 21 days, ADS for 14 days and representation for asthma for 21 days).

**Results:** PCR for viruses/atypical bacteria was positive in 81.7% of children (75.1% human rhinovirus, co-detection in 14.2%). *M. pneumoniae* and *C. pneumoniae* were rarely detected. The presence of micro-organisms had little impact on acute asthma or recovery outcomes. Children with atopy were significantly more likely to relapse and represent for medical care by day-14 (OR 1.11, 95%CI 1.00,1.23).

**Conclusions:** The presence of any viruses is associated with asthma exacerbations but does not appear to influence asthma recovery. In contrast, atopy is associated with asthma relapse. *M. pneumoniae* and *C. pneumoniae* are rare triggers of acute asthma in young children.

## INTRODUCTION

Acute asthma, one of the most common causes of childhood emergencies, is the subject of many publications based in Emergency Departments (EDs)[1, 2]. However, there is paucity of data on the recovery period as most studies have limited outcomes to length-of-stay, medications, hospital admission and relapse. Most children with asthma exacerbations are not hospitalised but many have respiratory morbidity lasting >2 weeks[3, 4].

It is likely that many factors govern the severity of acute asthma on presentation and the recovery period reflecting on-going morbidity in children. These factors include extrinsic determinants (e.g. access to service and socioeconomic influences[5]) and biological factors. Data on the latter are scarce in children; possible factors are the presence of viral infections[2, 6] and atopy[7, 8].

Viral infections are detected in up to 80% of children with asthma exacerbations[6]. Although the presence of atypical bacteria (*Mycoplasma*/*Chlamydophila)* has been associated with unstable asthma[9-11], few studies have evaluated the impact of respiratory viral and atypical bacteria detection on acute asthma severity or symptom resolution during the recovery period. One study[2] reported that viral infection symptoms were associated with poorer response to β_2-_agonists while another[12] reported that virus detection by PCR did not impact on acute severity or resolution of asthma quality of life (AQOL)[13] and diary scores (ADS)[14], although the findings were limited to non-hospitalised children.

Viruses in conjunction with allergens or atopic eczema likely increase the risk of hospitalisation[7, 15] or severe asthma[8]. Paediatric studies have described an association between allergic sensitization and/or exposure to inhalant allergens and risk for hospitalisation for acute asthma[16, 17]. However to our knowledge, no paediatric studies have examined the influence of atopy on asthma morbidity (i.e. the recovery) following acute exacerbations.

We evaluated the impact of respiratory viruses/atypical bacteria and atopy on the acute severity and clinical recovery in 244 children presenting to EDs with acute asthma (hospitalised and non-hospitalised children). We hypothesized that symptoms of asthma exacerbations are more severe and prolonged in children with a respiratory virus/atypical bacteria or atopy. Our secondary aim was to describe the point prevalence of various respiratory viruses and atypical bacteria.

## METHODS

### Subjects

Children aged 2-16 years who presented with an acute asthma exacerbation to the ED at 2 hospitals [Royal Children's Hospital (RCH, Brisbane), July 2009-December 2010 and Canberra Hospital (TCH), January 2010-June 2011] were recruited. Written informed consent was obtained from a parent/carer.

Asthma was defined as recurrent (>2) episodes of wheeze and/or dyspnoea with a clinical response (decreased respiratory rate and work of breathing) to β_2_-agonist, as diagnosed by a doctor unrelated to this study. Asthma exacerbation was defined as an acute deterioration of asthma control requiring treatment with >1 dose (>600µg via metered dose inhaler and spacer/>2.5mg nebulised) of salbutamol in an hour. Exclusion criteria for the study were presence of: an underlying respiratory disease (e.g. bronchiectasis), cerebral palsy/severe neurodevelopmental abnormality, immuno-compromised state, severe asthma (requiring continuous nebulised/intravenous salbutamol) or previously enrolled in the study. Children were managed by ED staff who were uninvolved in the study. The study was approved by the ethics committees of both hospitals.

### Study Protocol

Clinical history and examination were documented on a standardised data collection sheet, including questions specific for asthma (e.g. exacerbation frequency, medications) and for acute respiratory infection symptoms (ARI: runny nose, fever, sore throat, cough, irritability, tiredness). An ARI was considered present if ≥2 symptoms were present at enrolment[18]. Baseline asthma severity was determined using an Australian Functional Severity Scale (FSS) for paediatric asthma[19]. Severity of acute asthma on presentation was categorised according to Acute Asthma Score[20] and the Australian National Asthma Guidelines (NAC)[21]. Children were treated by doctors in accordance with the Australian NAC using a standardised protocol. A nasopharyngeal aspirate (NPA) was undertaken for PCR detection of viruses, *Chlamydophila* and *Mycoplasma* (supplement) and treating doctors were unaware of the results. Skin prick tests (SPT) to 6 environmental allergens (supplement) were also performed. Children were considered atopic if a wheal ≥3mm in diameter to any allergen (above negative control) developed. Eczema (in the last 12-months) was self-reported using the International Study of Asthma and Allergies in Childhood (ISAAC) questionnaire.

Baseline and weekly asthma quality of life questionnaires for parents (PACQLQ)[13] and validated daily ADS[14] were recorded for 21 and 14-days respectively. Follow-up phone calls occurred 24-48 hours after enrolment and on days-7, 14 and 21 where PACQLQ and adverse events including unscheduled representations to a health facility were recorded. End points were exacerbation of asthma requiring corticosteroids, admission into hospital and/or at Day-21 (whichever occurred first).

The outcomes were divided into (a) acute severity outcomes (NAC assessment, hospitalisation, FSS, acute asthma score, PACQLQ on presentation, ADS on presentation and length of hospitalisation) and (b) recovery outcomes (PACQLQ on days-7, 14 and 21, ADS on days-5, 7, 10 and 14 and representation for asthma on days-7, 14 and 21).

### Statistics

Data were first examined using normality plots. Mann-Whitney U test was used for 2 group comparisons of non-normal data and Chi squared test (or Fisher's exact test when appropriate) for categorical variables. Data for the association between "PCR-positive state" (presence of any virus/atypical bacterium in NPA) and atopy (SPT positivity) with measures of acute severity and recovery outcomes were first examined using univariate analyses. This was then followed by multivariate linear regression to examine the association between measures of acute asthma severity and recovery outcomes with "PCR-positive state" and atopy while considering potential contributors (age, gender, inhaled corticosteroids use, presence of smokers in the household, days unwell before presentation). Two-tailed p value of <0.05 was considered significant. SPSS v.23.0 was used for statistical calculation.

## RESULTS

The characteristics of the 244 children (mean age 5.5±SD 3.1 years) enrolled are presented in Table 1. Sixty-eight of 86 (79.1%) children who had eczema were atopic and 68/158 (43.0%) who were atopic had eczema. Ninety of 121 (74.4%) children who did not have eczema were atopic and 18/49 (36.7%) who were not atopic had eczema.

**Table 1.** Characteristics of Children Enrolled

|  | **Brisbane (RCH)** | **Canberra (TCH)** | **Total** |
| --- | --- | --- | --- |
| Children enrolled, n | 136 | 108 | 244 |
| Age (years), median (IQR) | 4.2 (3.3) | 5.5 (4.7) | 4.5 (3.5) |
| Gender, n (F:M) | 53:83 | 32:76 | 85:159 |
| ETS exposure n (%) | 37 (27.2) | 37 (34.3) | 74 (30.3) |
| On inhaled corticosteroids n (%) | 44 (32.4) | 45 (41.7) | 89 (36.5) |
| On leukotriene receptor antagonist n (%) | 7 (5.1) | 9 (8.3) | 16 (6.6) |
| History of eczema ever present n (%) | 74 (54.4) | 70 (64.8) | 144 (59.0) |
| History of current eczema in the last 12 months n (%) | 54 (40.3) | 52 (48.6) | 106 (44.0) |
| Diagnosed allergy n (%) | 44 (32.4) | 30 (27.8) | 74 (30.3) |
| Pet exposure n (%) | 82 (60.3) | 71 (65.7) | 153 (62.7) |
| Days unwell before presentation, median (IQR) | 4.0 (2.0) | 3.0 (2.0) | 3.0 (1.0) |
| Acute Respiratory Infection present n (%)* | 127 (94.1) | 91 (88.3) | 218 (91.6) |
| Positive skin prick test n (%) | 85 (76.6) | 75 (76.5) | 160 (76.6) |
| % SpO2 on presentation, median (IQR) | 95 (4.0) | 91 (4.0) | 93 (6.0) |
| Respiratory rate on presentation, median (IQR) | 40.0 (18.0) | 35.5 (16.0) | 40.0 (16.0) |
| Asthma severity score, mean (SD)[20] | 5.4 (1.9) | 5.3 (1.5) | 5.3 (1.8) |
| Hospitalised % | 61.8 | 91.7 | 75.0 |
| Representation for asthma within 7 days (n=129 RCH, n=103 TCH); n (%) ^ | 21 (16.3) | 4 (3.9) | 25 (10.8) |
| Representation for asthma within 14 days (n=126 RCH, n=100 TCH); n (%) ^ | 18 (14.3) | 6 (6.0) | 24 (10.6) |

*ETS exposure indicates presence of any smoker in the household; \*Acute Respiratory Infection considered present if ≥ 2 symptoms (runny nose, fever, sore throat, cough, irritability, tiredness) present at enrolment. Data specific for the presence of an ARI were incomplete in 6 children. ^Denominator different for the various time points as data was missing on follow-up.*

### Point prevalence of viruses, *M. pneumoniae* and *C. pneumoniae*

NPAs were obtained from 243 children (Brisbane=135, Canberra=108). All NPA samples from Canberra and 117 (86.7%) from Brisbane were assessed using the extended viral/atypical bacterial PCR panel (supplement). Of these, PCR for various viruses/atypical bacteria was positive in 184/225 (81.7%) children [Brisbane: n=104/117 (88.9%); Canberra: n=80/108 (74.1%)]. PCR-positive children were significantly (p=0.002) younger than PCR-negative children.

Human rhinovirus (HRV) was present in 169/225 (75.1%) children who had the extended panel performed [Brisbane: n=95/117 (81.2%); Canberra: n=74/108 (68.5%)]. Enterovirus D68 was detected in 10%. Other viruses detected included RSV=7, hMPV=4, adenovirus=7, human bocavirus (HBoV)=14, WU polyomavirus (WUPyV)=13 and KI polyomavirus (KIPyV)=4. *M. pneumoniae*=3 (2 had concurrent HRV); all 225 specimens were negative for *C. pneumoniae.* Co-detection of micro-organisms occurred in 14.2% of children. Twenty-eight children were positive for 2 micro-organisms and 4 positive for 3 micro-organisms. Thirteen of 225 samples (5.8%) tested were positive for WUPyV; 11 of these had ≥1 additional virus detected [most common co-detections were HRV (n=9) and HBoV (n=4)]. There was no significant difference in PCR positive state in children with atopy (79.2%) versus no atopy, compared to children with eczema (85.7%) versus no eczema (p=0.678 and 0.224 respectively).

### Relationship between the presence of viruses/atypical bacteria with acute severity and recovery

Univariate analysis demonstrated that acute severity outcomes did not vary in relation to the PCR positive state (Table 2). On multivariate regression (adjusting for age, gender, inhaled corticosteroids and presence of smokers) comparing PCR-positive and negative groups, the only marker that was significantly different between PCR-positive and negative groups was PACQLQ. PACQLQ on admission was significantly higher (i.e. better) in those with PCR-positive state (β=0.40, 95%CI 0.04,0.75, p=0.028). Regression analyses revealed that PCR-state had no significant influence on NAC assessment on presentation, hospitalisation, length of hospitalisation, FSS, acute asthma score or ADS on presentation (p range 0.300-0.963).

**Table 2.** Acute Asthma Severity Outcomes Based on NPA Results and Atopy (Univariate Analysis)

|  | Nasopharyngeal aspirate |  |  | Atopy |  |  |
| --- | --- | --- | --- | --- | --- | --- |
| **Acute data**<br>**(on presentation):** | **Positive**<br>**(n=184)** | **Negative**<br>**(n=41 )** | **p** | **Positive**<br>**(n=160)** | **Negative**<br>**(n=49)** | **p** |
| NAC (moderate); n (%) | 123 (66.5) | 23 (56.1) | 0.283 | 103 (64.4) | 31 (63.3) | 0.897 |
| Hospitalisation; n (%) | 139 (75.1) | 32 (78.0) | 0.841 | 124 (77.5) | 34 (69.4) | 0.258 |
| FSS; mean (SD) | 8.3 (4.2) | 8.6 (5.3) | 0.629 | 8.2 (4.5) | 8.8 (4.2) | 0.440 |
| *Median (IQR)* |  |  |  |  |  |  |
| Acute asthma score[20] | 5.0 (3.0) | 5.0 (2.0) | 0.677 | 5.0 (2.0) | 6.0 (3.0) | 0.255 |
| PACQLQ score[13] | 4.8 (1.5) | 4.5 (2.1) | 0.182 | 4.8 (1.6) | 4.6 (1.6) | 0.543 |
|  | n=102 | n=21 |  | n=94 | n=20 |  |
| Asthma diary score[14] (n=123<br>NPA, 114 Atopy)* | 3.5 (2.6) | 4.0 (1.8) | 0.202 | 3.8 (2.5) | 3.3 (2.4) | 0.641 |
|  | n=139 | n=32 |  | n=124 | n=34 |  |
| Length of hospitalisation<br>(hours)* | 41.0 (23.0) | 43.0 (23.0) | 0.897 | 43.0 (24.0) | 38.0 (16.0) | 0.572 |

*NAC =National Asthma Council initial assessment[21]; FSS = functional severity scale[19]; Atopy = positive allergen skin prick test to one or more allergens*
*NPA unavailable in 18 children; \*data incomplete for these outcomes.*

Analyses of recovery outcomes on univariate analysis (Table 3) show that ADS were significantly better on days-5 and 10 in those who were PCR-positive compared to the PCR-negative group. However, this difference disappeared at later time points. There was no significant difference between groups for any other recovery outcome.

**Table 3.** Asthma Recovery Outcomes Based on NPA Results and Atopy (Univariate Analysis)

|  | Nasopharyngeal aspirate |  |  |  | Atopy |  |  |
| --- | --- | --- | --- | --- | --- | --- | --- |
| Recovery data (on follow-up): | Positive | Negative | P |  | Present | Absent | P |
| *Median (IQR)* | n=170 | n=39 |  |  | n=149 | n=45 |  |
| PACQLQ day 7 [13]<br>(n=209 NPA, 194 Atopy) | 6.0 (1.7) | 6.0 (1.8) | 0.866 |  | 6.0 (1.5) | 5.2 (2.3) | **0.044** |
|  | n=164 | n=38 |  |  | n=144 | n=42 |  |
| PACQLQ day 14 (n=202<br>NPA, 186 Atopy) | 6.7 (0.9) | 6.7 (1.0) | 0.243 |  | 6.8 (0.8) | 6.5 (1.2) | 0.112 |
|  | n=164 | n=36 |  |  | n=140 | n=46 |  |
| PACQLQ day 21 (n=200<br>NPA, 186 Atopy) | 6.8 (0.8) | 6.7 (1.2) | 0.315 |  | 6.8 (0.6) | 6.7 (1.1) | 0.063 |
| Asthma diary score[14]<br>on: | n=101 | n=19 |  |  | n=93 | n=19 |  |
| Day 5 (n=120 NPA, 112<br>Atopy) | 1.5 (2.3) | 2.3 (2.3) | **0.012** |  | 1.5 (2.0) | 1.5 (2.8) | 0.818 |
|  | n=98 | n=19 |  |  | n=89 | n=19 |  |
| Day 7 (n=117 NPA, 108<br>Atopy) | 1.5 (1.8) | 1.8 (1.3) | 0.096 |  | 1.5 (1.8) | 1.3 (2.3) | 0.696 |
|  | n=95 | n=19 |  |  | n=87 | n=19 |  |
| Day 10 (n=114 NPA,<br>106 Atopy) | 0.3 (1.8) | 1.5 (1.0) | **0.045** |  | 1.0 (1.5) | 0.3 (1.5) | 0.429 |
|  | n=89 | n=18 |  |  | n=81 | n=19 |  |
| Day 14 (n=107 NPA,<br>100 Atopy) | 0.5 (1.5) | 1.5 (1.5) | 0.137 |  | 0.5 (1.5) | 0.3 (1.5) | 0.822 |
|  | n=175 | n=40 |  |  | n=150 | n=47 |  |
| Representation for<br>asthma within 7 days<br>(n=215 NPA, 197<br>Atopy); n (%) | 20 (11.4) | 3 (7.5) | 0.581 |  | 13 (8.7) | 7 (14.9) | 0.267 |
|  | n=169 | n=40 |  |  | n=147 | n=46 |  |
| Representation for<br>asthma within 14 days<br>(n=209 NPA, 193<br>Atopy); n (%) | 20 (11.8) | 2 (5.0) | 0.263 |  | 17 (11.6) | 1 (2.2) | 0.078 |
|  | n=164 | n=36 |  |  | n=140 | n=46 |  |
| Representation for<br>asthma within 21 days<br>(n=200 NPA, 186<br>Atopy); n (%) | 12 (7.3) | 2 (5.6) | 1.000 |  | 8 (5.7) | 4 (8.7) | 0.495 |

*Atopy indicates positive allergen skin prick test.*

Multivariate regression confirmed that the PCR-positive state when children presented to ED was associated with clinical scores that were generally better during recovery. ADS on day-5, but not at later times, were better in children with a PCR-positive state (β=;-0.71, 95%CI-1.36,-0.06, p=0.032). Similarly, PACQLQ on day-14 and 21 was significantly higher (better) in those with PCR-positive state (β=0.29, 95%CI 0.01,0.57, p=0.043 and β=0.35, 95% CI 0.05,0.64, p=0.022 respectively). On multivariate regression, PCR-state had no significant influence on other recovery outcomes (representation for asthma on days-7, 14 and 21) (p range 0.164-0.846).

### Effect of atopy on acute severity and recovery

On univariate (Table 2) and multivariate analyses, all the acute severity outcomes were not significantly associated with atopy (p range 0.368-0.998 for multivariate regression).

Children with atopy had significantly higher (better) PACQLQ on day-7 on univariate analysis (Table 3). However, this difference disappeared in later days. On multivariate regression, atopy was not significantly associated with PACQLQ on days-7, 14 or 21 (p range 0.082-0.414).

For the recovery period, atopy was not significantly associated with ADS on univariate analysis (Table 3). This remained the case also on multivariate analysis (p range 0.222-0.795). On multivariate regression, atopy was not significantly associated with other recovery outcomes (representation for asthma on days-7 and 21) (p=0.160 and 0.659 respectively) but was significantly associated with representation for asthma on day-14. Children with atopy were significantly more likely to relapse and represent for asthma deterioration by day-14 (OR 1.11, 95%CI 1.00,1.23, p=0.042).

### Relative impact of PCR-positive state, current eczema and atopy on asthma outcomes

Restricting this multi-regression analysis to major outcomes for each of the asthma phases (supplement), we found no significant influence on hospitalisation. However, PACQLQ on admission and day-14 were significantly higher (better) in those with PCR-positive state (β=0.48, 95%CI 0.10,0.85, p=0.013 and β=0.33, 95%CI 0.05,0.62, p=0.022 respectively). PCR-state was not significantly associated with PACQLQ on day-7. Atopy was not significantly associated with PACQLQ on admission and on day-14 but was associated with PACQLQ on day-7 with higher (better) scores in those with atopy (β=0.45, 95%CI 0.09,0.82, p=0.015). PCR-state, eczema and atopy were not significantly associated with ADS on day-10.

## DISCUSSION

We examined the factors associated with measures of acute severity and clinical recovery in 244 children with hospitalised and non-hospitalised asthma exacerbations. Using PCR for an extended panel of viruses/atypical bacteria, micro-organisms were detected in 81.7% of children. However, PCR-positivity had little impact on acute severity or recovery outcomes: PACQLQ was actually better on presentation and during recovery (day-14 and 21) in PCR-positive compared to PCR-negative patients, though the difference was small. The presence of atopy did not impact on any measure of acute asthma or recovery outcomes. However, children with atopy were significantly more likely to represent for asthma deterioration by day-14.

### Respiratory viruses/atypical bacteria and effect on acute severity and recovery

Our PCR-positive rate is similar to that described by Johnston et al[6] (80%) but higher than others of 63-64%[2, 18]. Our rate of PCR identification of micro-organisms is higher than our previous study (54%) of non-hospitalised children[12]. In addition, in this study we included additional respiratory viruses (polyomaviruses and HBoV). Like others[12, 22], we found that viruses were more likely to be present in younger children and that HRV was the most frequent virus identified[12, 18]. In our cohort, *M. pneumoniae* and *C. pneumoniae* did not seem important, unlike 1 study on acute wheeze[23], but similar to other studies on asthma[24].

There are only a few studies on the impact of viral detection on measures of acute severity and recovery. In our previous study[12] involving only non-hospitalised children, only 78 of the 201 children had an NPA performed. Our current larger study confirms that the presence of a viral respiratory illness had a modest influence on acute severity and recovery from an asthma exacerbation. Children who were PCR-positive had significantly better PACQLQ scores than PCR-negative children but the difference between groups (β of 0.29 and 0.35) was less than the minimal important difference of 0.5[25]. Nevertheless, this suggests (a postulate) that other extrinsic factors (e.g. traffic-related air pollution[26, 27]) may have triggered the asthma exacerbations of children with PCR-negative state, resulting in a longer duration of symptoms in the children who were PCR-negative.

### Association between atopy and acute severity and recovery

We considered it important to differentiate between eczema and atopy in light of recent studies[28]- 30]. While previous studies have demonstrated that 45-64% of patients with eczema are non-atopic, and children with non-atopic eczema have a lower risk of developing asthma than those with atopic eczema[28]-31], we found that 79.1% of our children with eczema were also atopic.

In a small case-control study (n=60 inpatients), Green[7] described that adults who were hospitalised were more likely to be sensitised (by skin prick test) and exposed to either mite, cat, or dog allergen than patients with stable asthma (37%) and inpatient controls (15%; p<0.001). Likewise we found that children with atopy were significantly more likely to represent to a doctor for relapse than those without (OR 1.11, 95%CI 1.00,1.23, p=0.042). However as a group, those with atopy had similar hospitalisation rates and scores in the recovery phase. While the acute representation may reflect parental effects, this is unlikely given that PACQOL was better in the atopic group (on univariate analysis). Xepapadaki et al's study[32] suggested that an increased rate of symptomatic cold and asthma episodes in atopic children was associated with considerable cumulative prolongation of airway hyper-responsiveness, which may help explain the role of atopy as a risk factor for asthma persistence.

One of our study’s novel aspects includes the focus on asthma recovery outcomes. This is important as the morbidity of asthma extends well beyond the immediate exacerbation phase[3]. We used patient-oriented and validated outcomes (PACQLQ and ADS). Patient-oriented outcomes are arguably as important as objective measures[33], which are limited in routine clinical care, especially in young children. We also examined the influence of viruses and atypical bacteria and atopy on acute severity and during the recovery period. This information is potentially important in identifying the children who are more likely to have an asthma relapse, with substantial burden placed on their parents/carers and families. Data could also aid in counselling parents of children with acute asthma regarding the potential length of symptoms and consequences.

There are several limitations to our study. Firstly, we did not examine for bacterial infection. It is possible that the children with bacterial infections may take longer to recover from an asthma exacerbation as bacterial infection has been shown to be important in acute wheeze[34]. Secondly, we limited our study to clinical matters and did not evaluate possible mechanisms underlying the higher risk of relapse in atopic children. While we may speculate that atopy might be associated with delayed resolution of airway inflammation, addressing this possibility would require further prospective studies. Thirdly, there were differences between these sites e.g. the hospitalisation and representation rate. Reasons for this are unknown but not unreasonable given that RCH/Brisbane is a tertiary hospital whereas TCH/Canberra is not.

We conclude that although asthma exacerbations are commonly associated with viruses, their presence does not impact on recovery. In addition, children with atopy are more likely to have an unscheduled doctor visit within 14-days. Also, *M. pneumoniae* and *C. pneumoniae* are rare triggers of acute asthma in young children.

## CONTRIBUTORS

LT contributed to the conception and design, acquisition of data, analysis and interpretation of data and writing of the manuscript. PVA, JA and MH contributed to the conception and design, supervision and revision of the manuscript. WHS and CYTW contributed to the acquisition of data. IMM and TPS contributed to the acquisition of data and revision of the manuscript. JWU contributed to the interpretation of data and revision of the manuscript. TN contributed to the analysis and interpretation of data and revision of the manuscript. ABC contributed to the conception and design, supervision, interpretation of data and revision of the manuscript. All authors approved the final manuscript.

## ACKNOWLEDGEMENT

A component of this manuscript has been presented at the Thoracic Society of Australia and New Zealand Annual Scientific Meeting in 2014.

## What is already known on this topic

Viral infections are detected in up to 80% of children with asthma exacerbations.

Viruses in conjunction with allergens or atopic eczema likely increase the risk of hospitalisation or severe asthma.

## What this study adds

This study of 244 children enrolled in 2 Australian centres found that the presence of atopy increased the risk of representation for asthma relapse.

The presence of viral detection had minimal impact on acute asthma severity or recovery.

